# Mind-wandering in Parkinson’s disease hallucinations reflects primary visual and default network coupling

**DOI:** 10.1101/347658

**Authors:** Ishan C. Walpola, Alana J. Muller, Julie M. Hall, Jessica R. Andrews-Hanna, Muireann Irish, Simon J. G. Lewis, James M. Shine, Claire O’Callaghan

## Abstract

A mismatch between top-down expectations and incoming sensory information is thought to be associated with hallucinations across a range of neuropsychiatric disorders. In Parkinson’s disease with visual hallucinations, abnormal activity within the default network, and its pattern of connectivity with early visual regions, has been identified as a potential pathological source of the internally generated expectations that override incoming sensory input. In the context of attention deficits and visual dysfunction, mental imagery and perceptual expectancies generated across the default network are suggested to exert excessive influence over earlier visual regions, leading to aberrant perceptual experiences. Whilst converging neuroimaging evidence has identified unconstrained default network activity in Parkinson’s disease with hallucinations, to date there has been a lack of behavioural evidence to confirm the consequences of an over-engaged default mode network – therefore the contributions it might make to hallucination phenomenology remain speculative. To address this, we administered a validated thought-sampling task to 38 Parkinson’s disease patients (18 with hallucinations; 20 without) and 40 controls, to test the hypothesis that individuals with hallucinations experience an increased frequency of mind-wandering – a form of spontaneous cognition strongly associated with mental imagery and default network activity. The neural correlates of mind-wandering frequency were examined in relation to resting-state functional connectivity. Our results showed that patients with hallucinations exhibited significantly higher mind-wandering frequencies compared to non-hallucinators, who in turn had reduced levels of mind-wandering relative to controls. Inter-network connectivity and seed-to-voxel analyses confirmed that increased mind-wandering in the hallucinating vs. non-hallucinating group was associated with greater coupling between the primary visual cortex and dorsal default network. Taken together, both elevated mind-wandering and increased default-visual network coupling emerged as a distinguishing feature of the hallucinatory phenotype. We propose that the finding of increased mind-wandering reflects unconstrained spontaneous thought and mental imagery, which in turn furnish the content of visual hallucinations. Our findings suggest that primary visual cortex to dorsal default network coupling may provide a neural substrate by which regions of the default network exert disproportionate influence over ongoing visual perception. These findings refine current models of visual hallucinations by identifying a specific cognitive phenomenon and neural substrate consistent with the top-down influences over perception that have been implicated in visual hallucinations.

## Introduction

Hallucinations that are predominantly visual in nature affect over 40% of patients in the early stages of Parkinson’s disease, and upwards of 80% as the disease progresses (Ffytche *et al.*, 2017). Parkinson’s disease visual hallucinations have been associated with increased activity and connectivity in the default network, both in the resting state (Yao *et al.*, 2014; Franciotti *et al.*, 2015) and during recorded visual misperceptions (Shine *et al.*, 2015b). The default network is situated most distantly from primary sensory networks along a hierarchical gradient of cortical connectivity (Margulies *et al.*, 2016; Huntenburg *et al.*, 2018). As such, it is a key candidate for generating certain high-level predictions to prepare earlier visual regions for incoming sensory information, thereby increasing perceptual sensitivity and efficiency (de Lange *et al.*, 2018).

Several models of Parkinson’s disease visual hallucinations have posited that in the context of poor quality visual input – due to attentional impairments, visual deficits and retinal pathology – higher-order regions involved in the generation of mental imagery and perceptual expectancies exert excessive influence upon perception (Collerton *et al.*, 2005; Diederich *et al.*, 2005; Shine *et al.*, 2014). An instantiation of such models is that an over-active or unconstrained default network dominates the perceptual processes in a top-down manner, supplying perceptual predictions in the form of internally generated imagery that overrides incoming sensory information (Shine *et al.*, 2014; Powers *et al.*, 2016; O’Callaghan *et al.*, 2017b). This may be further exacerbated in Parkinson’s disease due to the numerous sources of visual and attentional dysfunction that render incoming sensory information less reliable (Weil *et al.*, 2016). Such an imbalance between top-down influences and incoming sensory input can be interpreted in a Bayesian predictive coding framework, in which increased precision (i.e., relative weighting) is consistently afforded to prior beliefs (i.e., top-down expectations or predictions), such that they override incoming sensory evidence and dominate the ultimate percept (Friston, 2005; Fletcher and Frith, 2009; Adams *et al.*, 2013). Attentional network dysfunction – specifically involving the dorsal attention network – has been established in Parkinson’s disease with visual hallucinations, and may reflect an inability to dynamically modulate precision at the mesoscale of brain function (Shine *et al.*, 2015a; 2015b; Hall *et al.*, 2019). This is in keeping with a recent finding that in Parkinson’s disease with visual hallucinations, accumulation of sensory evidence is slow and inefficient – and is therefore less informative – which may cause it to be down-weighted in favour of relatively preserved perceptual priors (O’Callaghan *et al.*, 2017a).

The hierarchical nature of the brain’s visual processing system supports the reciprocal feed-forward / feed-back flow of information between early visual and higher-order transmodal regions across the cerebral cortex (Gilbert and Li, 2013). More specifically, regions within the default network have been identified as sources of top-down influence over visual perception. These regions include the orbitofrontal and medial prefrontal cortices, which use early low spatial frequency information to generate expectations that constrain ongoing visual processing (Bar, 2003; Bar *et al.*, 2006; Summerfield *et al.*, 2006; Kveraga *et al.*, 2007; Chaumon *et al.*, 2014); hippocampal pattern completion mechanisms that supply memory-based expectations to the visual cortex (Hindy *et al.*, 2016); parahippocampal and retrosplenial cortices supporting the rapid activation of contextual associations during visual processing (Kveraga *et al.*, 2011; Aminoff *et al.*, 2013); and, distinct populations of neurons in the inferior temporal cortex that encode predictions and prediction errors in relation to incoming visual input (Bell *et al.*, 2016; Kok, 2016). The temporal properties of supramodal regions such as the default network are also consistent with the unfolding of high-level predictions over longer timescales (Margulies *et al.*, 2016; Baldassano *et al.*, 2017; Weilnhammer *et al.*, 2018), which fits well with the complex and temporally extended hallucinations that occur in Parkinson’s disease (Ffytche *et al.*, 2017). Yet, despite a number of established routes by which the default network may influence visual perception, and evidence of unconstrained default network activity in Parkinson’s disease visual hallucinations, we know very little about the behavioural consequences of an over-engaged default network and how this might contribute to hallucinations.

In keeping with its role as a source of top-down influence over visual perception, the default network is implicated in many cognitive processes relevant for generating expectations about the sensory environment, including mental imagery and scene construction, autobiographical memory, prospection, and retrieval of contextual associations (Buckner *et al.*, 2008; Spreng *et al.*, 2009; Andrews-Hanna *et al.*, 2010; Kveraga *et al.*, 2011; Schacter *et al.*, 2012). One common experience that draws upon the aforementioned cognitive processes, and is strongly linked to activity in the default network, is mind-wandering. Mind-wandering is often characterised as thoughts that are decoupled from the immediate perceptual environment and unrelated to ongoing task demands (Smallwood and Schooler, 2015; Seli *et al.*, 2018). Recent frameworks further emphasise that mind-wandering is a mental state that arises spontaneously, in which thoughts are unguided and unconstrained (Christoff *et al.*, 2016; Irving, 2016).

There is a striking similarity between certain core characteristics of both mind-wandering and visual hallucinations: both are transient forms of spontaneous cognition, relatively unconstrained by sensory input, and are underpinned by dynamic shifts in the interactions both within and between similar large-scale brain networks (Christoff *et al.*, 2016; Collerton *et al.*, 2016; Zabelina and Andrews-Hanna, 2016; Kucyi, 2017). Given the shared phenomenology and evidence for an overlapping neural basis, we predicted that clinical subgroups with visual hallucinations would also show changes in their propensity for mind-wandering. While such definitive studies have not been conducted to date, in patients with schizophrenia higher frequencies of task-unrelated thought have been observed, correlating with the severity of their positive symptoms, including hallucinations (Shin *et al.*, 2015). Additionally, previous work in Parkinson’s disease has demonstrated that individuals with visual hallucinations exhibit stronger mental imagery – a prominent feature of mind-wandering – during a binocular rivalry paradigm (Shine *et al.*, 2015a). Taken together, these findings suggest that increased mind-wandering may provide a cognitive correlate for the predisposition to experience hallucinatory phenomena across disease states.

To address the question of whether mind-wandering is related to hallucinations in Parkinson’s disease, the present study utilised a validated thought sampling task designed for clinical populations with cognitive impairment (O’Callaghan *et al.*, 2015). Using this task, we measured mind-wandering frequencies in Parkinson’s disease patients with and without visual hallucinations, and healthy controls. To explore the neural correlates of mind-wandering frequency, we used network-level and seed-to-voxel analysis of resting-state functional magnetic resonance imaging. We were primarily interested in between-group differences in the association between mind-wandering frequency and network interactions in patients, to determine the features uniquely related to hallucination predisposition. Based on previous work, we predicted that connectivity between the default network and visual areas would be related to elevated mind-wandering in patients with visual hallucinations. Contrasts with healthy controls were of secondary interest and were used to inform differential patterns of brain connectivity and behaviour potentially indicative of differential disease trajectories in the Parkinson’s disease groups. Overall, our aim was to investigate whether elevated mind-wandering and its associated neural correlates may be identifiable traits in a population prone to visual hallucinations.

## Methods and Materials

### Case selection

Thirty-eight individuals with Parkinson’s disease were recruited from the Parkinson’s disease research clinic, University of Sydney, Australia. Question two of the MDS-UPDRS (Goetz *et al.*, 2008) was used to identify visual hallucinations (i.e., “Over the past week have you seen, heard, smelled or felt things that were not really there? If yes, examiner asks the patient or caregiver to elaborate and probes for information”). If an individual scored ≥1 on this item and if their subsequent description was consistent with visual hallucinatory phenomena, including minor (passage or illusions) or complex hallucinations, they were included in the hallucinating group. This resulted in 18 patients with hallucinations and 20 without hallucinations.

All individuals with Parkinson’s disease satisfied the United Kingdom Parkinson’s Disease Society Brain Bank criteria and did not meet criteria for dementia, scoring above the recommended cut-off of ≥ 26 on the Montreal Cognitive Assessment (MoCA) (Dalrymple-Alford *et al.*, 2010). Motor severity was determined by the Hoehn and Yahr Scale and the unified Parkinson’s disease rating scale (MDS UPDRS-III) (Goetz *et al.*, 2008). Mood was assessed via the self-reported Beck Depression Inventory-II (BDI-II) (Beck *et al.*, 1996). General neuropsychological measures assessed working memory (backwards digit-span), attentional set-shifting (Trail Making Test, Part B minus Part A), and memory (story retention on the Logical Memory component of the Wechsler Memory Scale). All clinical and neuropsychological assessments, as well as neuroimaging, were performed with participants on their regular antiparkinsonian medication. Dopaminergic dose equivalence (DDE) scores were calculated, and no participants were taking antipsychotic medication or cholinesterase inhibitors. All individuals with Parkinson’s disease underwent neuroimaging.

Forty age- and education-matched healthy controls were included to provide a large normative dataset for the mind-wandering task (a subset of 20 of these controls also underwent neuroimaging). Controls were screened for a history of neurological or psychiatric disorders. The study was approved by the local Ethics Committees and all participants provided written informed consent in accordance with the Declaration of Helsinki. See Table 1 for demographic details and clinical characteristics.

**Table 1.** Mean (standard deviation) values for demographics, clinical characteristics and background neuropsychology.

| **Demographics, clinical characteristics & general neuropsychology** | **Controls** | **PD+VH** | **PD-VH** | ***p* value** |
| --- | --- | --- | --- | --- |
| **N** | 40 | 18 | 20 | - |
| **Sex (M:F)** | 21:19 | 14:4 | 17:3 | - |
| **Age** | 66.3 (6.2) | 67.5 (6.7) | 63.7 (6.6) | n.s. |
| **Education** | 14.7 (2.4) | 13.3 (3.3) | 14.6 (2.4) | n.s. |
| **MoCA** | - | 27.9 (1.3) | 27.9 (1.1) | n.s. |
| **Duration (yrs diagnosed)** | - | 7.6 (5.0) | 5.7 (3.2) | n.s. |
| **DDE (mg/day)** | - | 832.8 (395.7) | 691.5 (440.5) | n.s. |
| **Hoehn & Yahr stage** | - | 2.2 (.57) | 2.1 (.36) | n.s. |
| **UPDRS III** | - | 33.4 (15.9) | 28.4 (13.4) | n.s. |
| **BDI-II** | - | 11.6 (10.2) | 8.1 (5.9) | n.s. |
| ***Neuropsychology*** |  |  |  |  |
| **TMT B-A (seconds)** | - | 76.6 (57.3) | 35.2 (22.7) | ** |
| **Digit span backward** | - | 7.0 (1.6) | 6.6 (2.3) | n.s. |
| **Logical memory % retention** |  | 75.6 (17.7) | 87.1 (10.9) | * |

n.s. = non significant; \*\* = $p < .01$ ; \* = $p < .05$ . Significance tests refer to two-sided Welch's Independent samples t-tests or one-way ANOVAs with Tukey post hoc tests. PD-VH = non-hallucinators; PD+VH = hallucinators; MoCA = Montreal Cognitive Assessment; DDE = Dopaminergic dose equivalence; BDI-II = Beck Depression Inventory-II; TMT B-A = Trail Making Test part B minus part A.

### Mind-wandering experimental task & scoring procedures

The thought-sampling task was designed for use in patient populations with cognitive impairment and has previously been validated in older adults (O’Callaghan *et al.*, 2015). The task involved 9 trials. In each trial, a 2-dimensional coloured shape (e.g., blue square, yellow circle, etc.) was presented on the screen for varying durations (Short: ≤20 s, Medium: 30-60 s, Long: ≥90 s). At the outset, participants were told that they would be shown a series of shapes and to just relax and continue looking at the shape. Immediately after each shape was presented, the participant was prompted to describe aloud what they were thinking about during the presentation of the shape stimulus.

Participants’ reported thoughts were scored on a continuum, ranging from Level 1 to Level 4. Level 1 represents stimulus-bound/impoverished thought, including thinking about the stimulus, e.g., “a blue square,” or describing thinking of “nothing”. In contrast, Level 4 responses bear no obvious relationship to the stimulus, the task at hand, or the immediate testing environment, indicating thought content that is stimulus-independent and task-unrelated – a class of cognition often referred to as mind-wandering (Smallwood and Schooler, 2015). Examples of mind-wandering include, “I thought about the people I saw today and how we chatted with them outside the unit”; “I thought of a sailing boat in the Greek Islands”. Levels 2 and 3 represent intermediary responses, which do not qualify as fully-fledged instances of mind-wandering, as they still bear a discernible relationship to the presented stimulus or immediate environment. These levels capture the transition from stimulus-related to increasingly stimulus-independent responses. See Figure 1 and see Supplementary Material for a detailed description of the scoring levels and example responses.

**Figure 1.**
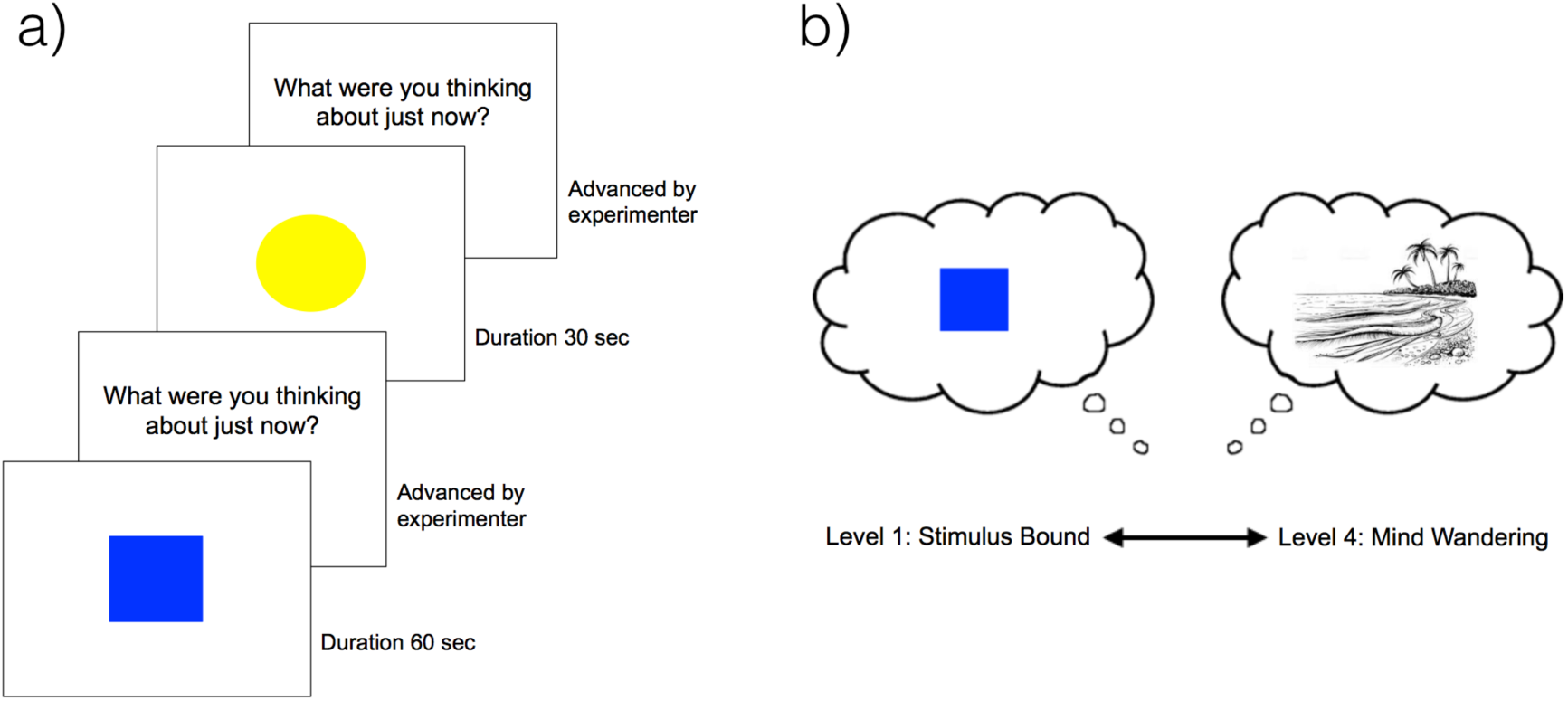
Task structure and schematic of scoring system. a) An example of two trials in the thought sampling task; b) Responses were scored from 1-4, with Level 1 consisting of a stimulus-bound response, such as reporting thoughts about the displayed shape, and Level 4 corresponding to thoughts completely unrelated to the task or immediate environment.

The final score awarded for each trial was the highest level achieved on that trial, ranging from 1-4. Instances of each scoring level achieved were counted across the 9 trials and transformed into a total percentage for each level during the task (i.e., total instances of levels 1, 2, 3 or 4 divided by 9 multiplied by 100). The primary analysis focused on differences in mind-wandering frequency between Parkinson’s disease with hallucinations, Parkinson’s disease without hallucinations, and controls. Therefore, the proportion of Level 4 responses – referred to as the mind-wandering frequency – was compared across the groups and used as a covariate in the neuroimaging analyses. Secondary behavioural analyses were conducted to determine overall performance of the three groups on the task. This involved comparing the proportion of responses that each group achieved across the four scoring levels of the task.

### Statistical analysis

Analyses were performed in R version 3.3.1 (http://www.r-project.org/). For performance on the mind-wandering task, homogeneity of variance was verified using Levene’s test and values were checked for normality by inspection of normal Q-Q plots and the Shapiro-Wilk test. To reduce skew in the data, a square root transformation was applied to the mind-wandering scores. Scores were then analysed using mixed-effects ANOVAs, implemented in the “lme4” package (Bates *et al.*, 2014). Where appropriate, group, score level and trial duration were specified as fixed effects, and subject was entered as a random effect. Where a factor only had three levels, post hoc analysis of significant main effects was performed using the Fisher’s Least Significant Difference (LSD) procedure (Cardinal and Aitken, 2013). Where factors had more than three levels, main effects were analysed using post-hoc t-tests with the Sidak correction for multiple comparisons. In these cases, post-hoc analyses of interactions were conducted using separate univariate ANOVAs to establish simple effects. Behavioural analysis scripts are available at https://github.com/claireocallaghan/MindWandering_PD_VH.

### Imaging acquisition

The 38 individuals with Parkinson’s disease and a subset of 20 controls underwent magnetic resonance imaging (MRI) to acquire T1-weighted structural images and resting-state blood-oxygenation level dependent (BOLD) functional scans. Imaging was conducted on a 3T MRI (General Electric, Milwaukee, USA). Whole-brain three dimensional T1-weighted sequences were acquired as follows: coronal orientation, matrix 256 x 256, 200 slices, 1 x 1 mm^2^ in-plane resolution, slice thickness 1 mm, TE/TR = 2.6/5.8 ms. T2*-weighted echo planar functional images were acquired in interleaved order with repetition time (TR) = 3 s, echo time (TE) = 32 ms, flip angle 900, 32 axial slices covering the whole brain, field of view (FOV) = 220 mm, interslice gap = 0.4 mm, and raw voxel size = 3.9 x 3.9 x 4 mm thick. Resting state scan acquisition lasted 7 minutes. During the resting-state scan, patients were instructed to lie awake with their eyes closed.

### Resting state fMRI preprocessing and motion correction

Functional magnetic resonance imaging (fMRI) pre-processing and analysis was performed using Statistical Parametric Mapping software (SPM12, Wellcome Trust Centre for Neuroimaging, London, UK, http://www.fil.ion.ucl.ac.uk/spm/software/). We used a standard pre-processing pipeline that included slice-timing correction, rigid body realignment, spatial smoothing with a Gaussian kernel (FWHM) of 6mm, and registration of the anatomical scans to the Montreal Neurological Institute standard brain space. Pre-processed images were imported into CONN: The Functional Connectivity toolbox (https://www.nitrc.org/projects/conn) in MATLAB for all functional connectivity analyses.

To compensate for motion-related artefacts we performed the “scrubbing” procedure in an effort to effectively remove time points with excessive head motion (i.e., framewise displacement in x, y, or z direction > 2 mm from the previous frame; global intensity > 9 standard deviations from mean image intensity of the entire resting state scan) (Power *et al.*, 2012). This approach (Artefact Detection Tools; https://www.nitrc.org/projects/artefact-detect/) effectively removes outlier frames by including them as dummy-coded regressors during the de-noising procedure, so as to avoid discontinuities in the time-series. We also tested for significant differences in maximum motion and number of frames scrubbed between the Parkinson’s disease groups. No significant differences were found in maximum framewise displacement (*p* = 0.50), maximum frames scrubbed (*p* = 0.25), mean framewise displacement (*p* = 0.59), or mean number of frames scrubbed (*p* = 0.25). Other noise sources in the BOLD signal (i.e., from white matter and cerebrospinal fluid) were corrected for by using a principle component-based ‘aCompCor’ method (Whitfield-Gabrieli and Nieto-Castañón, 2012). We applied a band-pass filter (0.008 - 0.09 Hz) to limit the effect of low-frequency drift and high-frequency noise on the BOLD signal time-series.

### Relationship between inter-network functional connectivity and mind-wandering frequency

To assess differences in the association between mind-wandering frequency and inter-network functional connectivity across the two patient groups and controls, we used 12 individual intrinsic connectivity network regions of interest, which were defined on a functional basis (Shirer *et al.*, 2012). The networks included in our analysis are shown in Figure 3a. We did not include the auditory or language networks, as the auditory network is not implicated in network-based models of visual hallucinations and, as others have noted, there is substantial overlap between the language network and aspects of the default/limbic networks (Zabelina and Andrews-Hanna, 2016). We corrected for multiple comparisons by using a False Discovery Rate (FDR) threshold of *p* < 0.05.

Based on the results from the above analysis, we performed a post-hoc seed-to-voxel functional connectivity analysis. This was to investigate whether the between-group differences (i.e., Parkinson’s disease hallucinators vs. non-hallucinators and controls vs. non-hallucinators) that were identified in the association between mind-wandering and primary visual network (V1) functional connectivity extended beyond the dorsal default network. We calculated a correlation between the average filtered BOLD signal in V1 and all other voxels in the brain for each group. Then, a between-group contrast of the regression coefficient, capturing the association between mind-wandering frequency and seed-to-voxel functional connectivity, was carried out with a height threshold of *p* < 0.001 and a cluster-size threshold of *p* < 0.05, corrected using FDR. Seed-to-voxel statistical tests for this post-hoc analysis were one-sided, as we were specifically interested in the spatial boundaries of the directional V1 connectivity demonstrated in the prior inter-network analysis.

## Results

### Demographics, clinical characteristics and background neuropsychology

Parkinson’s disease and control groups were matched for age and education. The Parkinson’s disease groups were matched on all demographic variables; however, the hallucinating group performed worse than the non-hallucinators on measures of attentional set-shifting and memory retention, although working memory performance did not differ significantly between the groups. See Table 1 and see Supplementary Material for detailed results.

### Hallucinators had a higher frequency of mind-wandering compared to non-hallucinators

As shown in Figure 2a, mind-wandering occurred significantly more frequently in hallucinators than non-hallucinators. There was a main effect of group [F(2,75) = 5.34, *p* < 0.01], and post-hoc comparisons revealed that controls and Parkinson’s disease with hallucinations did not differ, but both groups exhibited higher frequencies of mind-wandering than the non-hallucinating group (Fisher’s LSD, *p* < 0.05).

**Figure 2.**
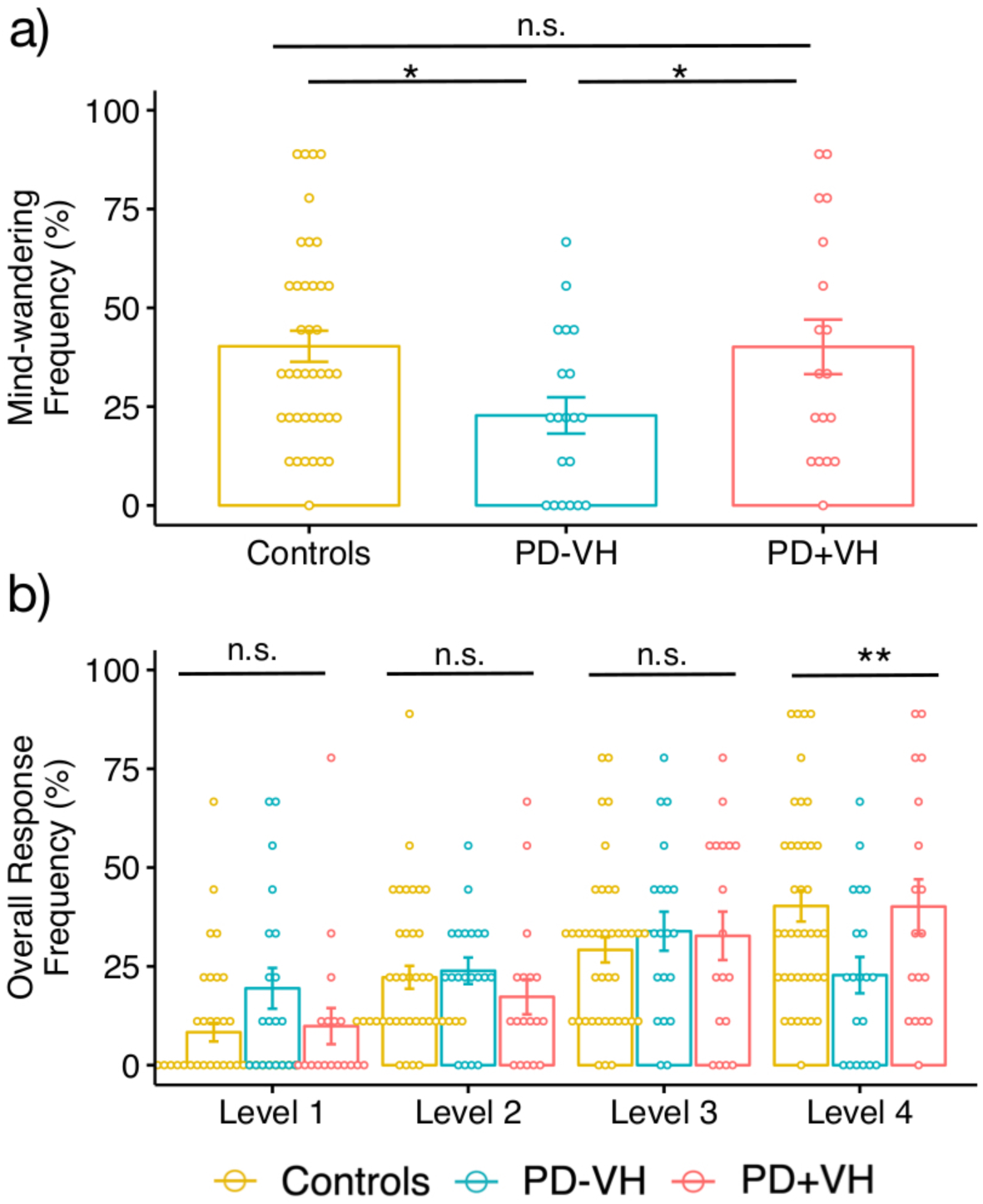
Performance on the mind-wandering task. a) Mind-wandering frequency. Parkinson’s disease with hallucinations and controls both exhibited higher frequencies of mind-wandering (i.e., had significantly more Level 4 responses on the thought sampling task) compared to non-hallucinators; b) Frequency of responses at each of the four scoring levels. Group differences were only found for Level 4 responses (i.e., mind-wandering frequency). Frequencies of responses across the three groups were not significantly different for Levels 1, 2 and 3. Error bars show standard error of the mean; PD-VH = non-hallucinators; PD+VH = hallucinators; n.s. = not significant; * = *p* < 0.05; ** = *p* < .01.

### Overall performance on the mind-wandering task revealed that significant group differences emerged exclusively for mind-wandering responses

Figure 2b shows the frequency of responses across all scoring levels, ranging from Level 1 (stimulus-bound thought) to Level 4 (mind-wandering). No significant main effect of group was evident [F(2,300) = 0.28, *p* = 0.76], however, a significant main effect of response level [F(3,300) = 28.57, *p* < 0.0001] was observed. Post hoc t-tests with Sidak correction showed that, regardless of group, Level 1 was the least frequent response, relative to Levels 2, 3 and 4 (*p* values < 0.0001), and higher frequencies of Level 4 were obtained relative to Level 2 (*p* < 0.01). The other Levels did not differ significantly from each other (*p* values > 0.05).

The Level x Group interaction was significant [F(6,300) = 3.06, *p* < 0.01]. We followed this interaction with tests of simple effects to determine whether the groups differed in their response frequencies at any Level apart from Level 4, which we showed in our focused analysis above. Follow-up tests of simple effects revealed the frequency of responses across the groups did not differ significantly for Level 1 [F(2,75) = 2.58, *p* = 0.08], Level 2 [F(2,75) = 1.16, *p* = 0.32] or Level 3 [F(2,75) = 0.20, *p* = 0.82]. The groups only differed significantly in their Level 4 responses (i.e., % mind-wandering frequency) [F(2,75) = 5.34, *p* < 0.01]. In addition, all groups showed an increased tendency towards mind-wandering on longer trials (See Supplementary material and Figure S1 for analyses and results), replicating previous studies using the task (Geffen *et al.*, 2017; O’Callaghan *et al.*, 2019).

### A stronger association was identified between primary visual-dorsal default inter-network coupling and mind-wandering frequency in Parkinson’s disease hallucinators vs. non-hallucinators

Results of the inter-network coupling analysis were consistent with our prediction of a stronger association between primary visual network and dorsal default network coupling and the degree of mind-wandering in patients with visual hallucinations. Of all network pairs (Figure 3a), only primary visual-dorsal default network functional connectivity (i.e., primary visual-dorsal default network coupling) and its association with mind-wandering frequency differed significantly between patient groups (*p* < 0.05, FDR; Figure 3b). Parkinson’s disease hallucinators had a significantly stronger positive association between mind-wandering frequency and primary visual-dorsal default network coupling compared with non-hallucinators. Controls also exhibited a stronger positive association between primary visual-dorsal default network coupling and mind-wandering frequency relative to Parkinson’s disease non-hallucinators, but this did not survive correction for contrasts among all network pairs. Only at a more lenient correction threshold – correcting for the number of contrasts between primary visual and all other networks (rather than all comparisons made) – was the association between controls and non-hallucinators statistically different (*p* < 0.05, FDR; Figure 3b). As this latter result only survived a more liberal correction for multiple comparisons, we emphasise caution in its interpretation. Statistically significant differences did not emerge between Parkinson’s disease hallucinators and controls.

**Figure 3.**
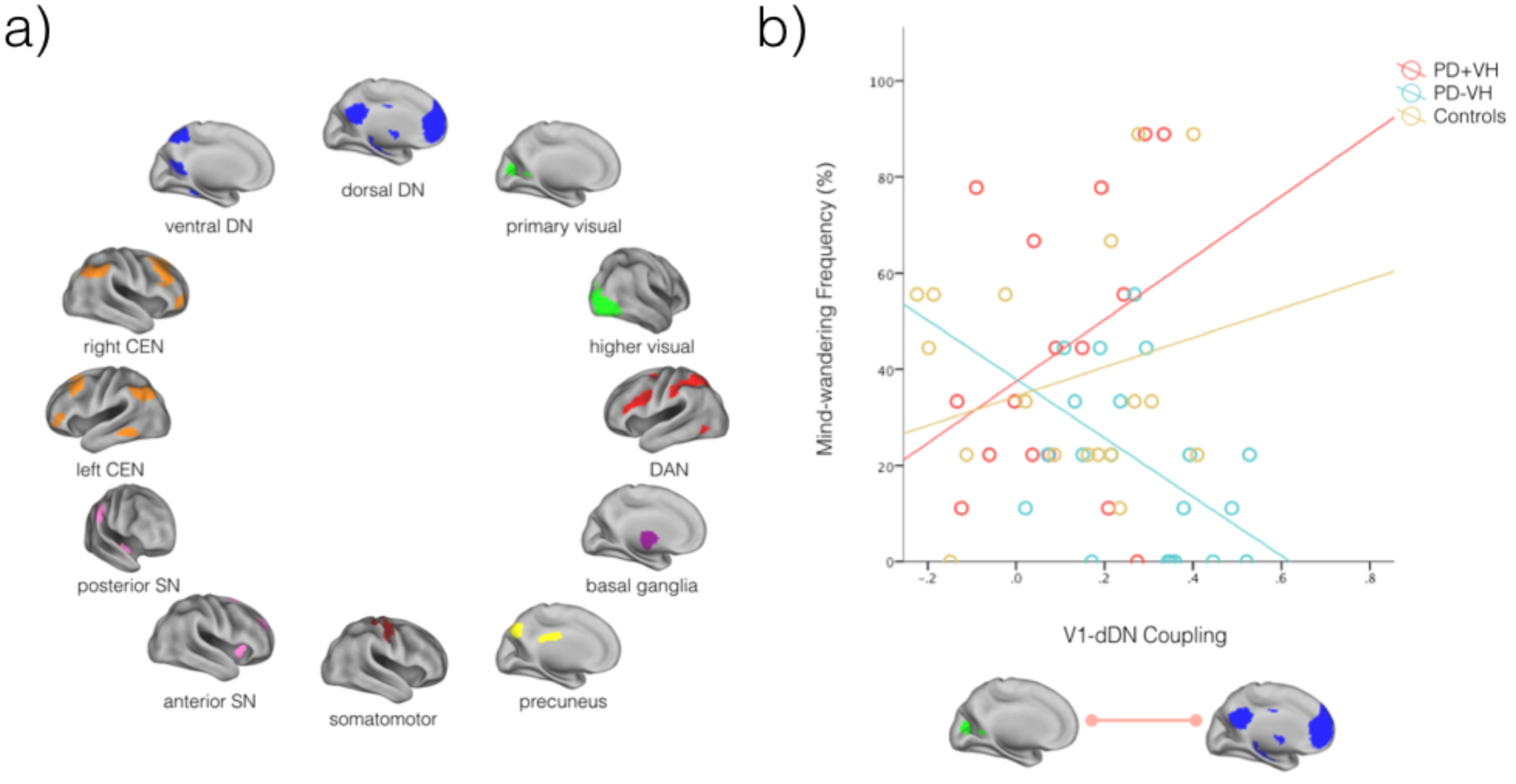
Association between mind-wandering frequency and inter-network coupling. a) Stanford atlas networks included in the analysis examining the association between inter-network functional connectivity and mind-wandering frequency; b) Individuals with visual hallucinations (PD+VH) had a significantly stronger positive association between mind-wandering frequency and V1-dDN coupling, compared to those without hallucinations (PD-VH). dDN = dorsal default network; vDN = ventral default network; CEN = central executive network; SN = salience network; DAN = dorsal attention network.

### Follow-up seed-to-voxel functional connectivity with a primary visual network seed

When the primary visual network was used as a seed, the between-group difference in the association between primary visual network coupling and mind-wandering frequency included brain regions both within and beyond the dorsal default network. Relative to the non-hallucinating group, hallucinators displayed a significantly stronger association between mind-wandering frequency and connectivity of the primary visual network seed to dorsal default network regions (posterior cingulate cortex, medial prefrontal cortex and left inferior parietal lobule), the inferior frontal gyrus, orbitofrontal cortex and high-level visual regions (fusiform gyrus/inferior temporal gyrus). See Figure 4 and Table 2. The same analysis contrasting controls and Parkinson’s disease non-hallucinators revealed similar core dorsal default network regions (posterior cingulate cortex and medial prefrontal cortex) as well as the orbitofrontal cortex. However, instead of unilateral inferior parietal lobule, changes were found with bilateral angular gyrus connectivity to primary visual cortex. Furthermore, the fusiform gyrus did not display altered primary visual cortex connectivity in this contrast (See Figure S2 and Table S1).

**Table 2.**
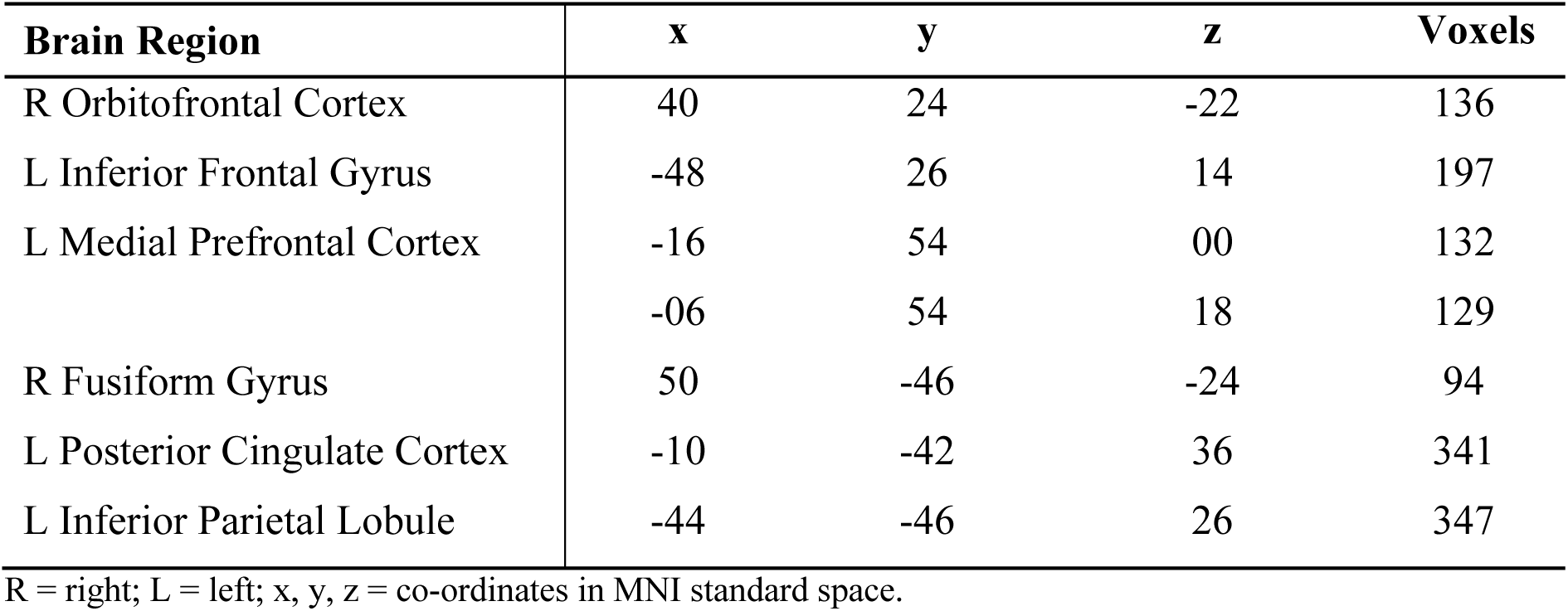
Peak coordinates from seed-to-voxel connectivity associated with mind-wandering frequency in Parkinson’s disease hallucinators vs. non-hallucinators

| Brain Region | x | y | z | Voxels |
| --- | --- | --- | --- | --- |
| R Orbitofrontal Cortex | 40 | 24 | -22 | 136 |
| L Inferior Frontal Gyrus | -48 | 26 | 14 | 197 |
| L Medial Prefrontal Cortex | -16 | 54 | 00 | 132 |
|  | -06 | 54 | 18 | 129 |
| R Fusiform Gyrus | 50 | -46 | -24 | 94 |
| L Posterior Cingulate Cortex | -10 | -42 | 36 | 341 |
| L Inferior Parietal Lobule | -44 | -46 | 26 | 347 |

R = right; L = left; x, y, z = co-ordinates in MNI standard space.

**Figure 4.**
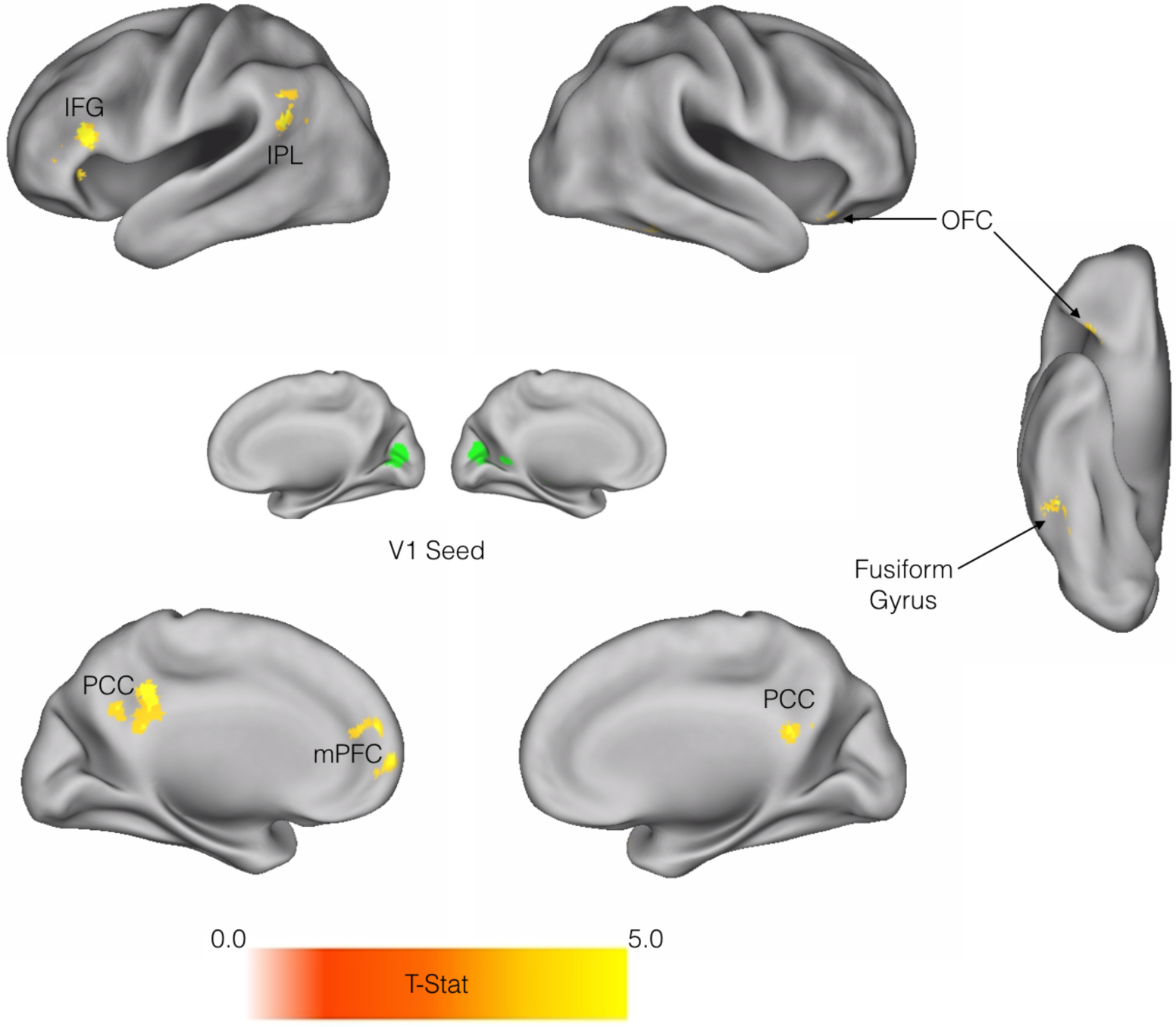
Seed-to-voxel connectivity associated with mind-wandering frequency, between V1 seed and whole brain in Parkinson’s disease hallucinators vs. non-hallucinators. Individuals with hallucinations had a significantly stronger association between mind-wandering frequency and connectivity between V1 and areas of the dorsal default network (PCC, mPFC and IPL), orbitofrontal cortex, inferior frontal gyrus, and high-level visual regions (fusiform gyrus/inferior temporal gyrus), relative to non-hallucinating individuals. PCC = posterior cingulate cortex; mPFC = medial prefrontal cortex; IPL = inferior parietal lobule; OFC = orbitofrontal cortex; IFG = inferior frontal gyrus.

## Discussion

Our results provide empirical evidence of a link between mind-wandering and hallucinations, revealing that Parkinson’s disease patients with visual hallucinations exhibit increased mind-wandering relative to non-hallucinating patients. Elevated mind-wandering may therefore represent a cognitive correlate of the excessive top-down influence upon perception that has previously been hypothesised in Parkinson’s disease visual hallucinations. Our resting state analysis revealed a route by which mind-wandering may impact upon early visual processing, as mind-wandering frequency was associated with stronger coupling between the primary visual and dorsal default networks in the Parkinson’s disease hallucinators. Together, these findings uncover trait characteristics: increased mind-wandering frequency related to default-visual network coupling, which may predispose the Parkinsonian brain to hallucinate.

Behavioural analysis of the thought-sampling task revealed that mind-wandering frequency was significantly higher in the hallucinating patient group compared to the non-hallucinating patient group; no significant differences were found between the hallucinating patient group and controls. In contrast, the frequency of mind-wandering was lower in non-hallucinating patients relative to controls, replicating findings from an independent Parkinson’s disease cohort tested on the same task (Geffen *et al.*, 2017). Preserved global cognitive function does not account for the higher levels of mind-wandering in Parkinson’s disease hallucinators, as their cognitive abilities were either similar to the non-hallucinators, or mildly reduced.

These findings suggest that reductions in mind-wandering may be common in Parkinson’s disease without hallucinations, consistent with observations in ageing (Jackson and Balota, 2012; McVay *et al.*, 2013; Irish *et al.*, 2018) and dementia populations including Alzheimer’s disease and frontotemporal dementia (Gyurkovics *et al.*, 2018; O’Callaghan *et al.*, 2019). By contrast, relatively preserved mind-wandering may be a specific feature of the hallucinating phenotype. This is consistent with theoretical models of Parkinson’s disease hallucinations that specify excessive influence of higher order visual regions over ongoing perception (Collerton *et al.*, 2005; Diederich *et al.*, 2005; Shine *et al.*, 2014). In Parkinson’s disease hallucinations, this excess of top-down influence is suggested to occur in the context of compromised primary visual and attentional systems, resulting in poor quality sensory evidence accumulation (Shine *et al.*, 2014; O’Callaghan *et al.*, 2017a). More broadly, this account is consistent with the idea that relatively strong prior beliefs are implicated in the genesis of hallucinations across modalities and across disorders (Corlett *et al.*, 2018).

Our correlations with imaging reveal a route by which overly strong prior beliefs or internal imagery might influence the early perceptual process. The primary finding was a significantly increased association between mind-wandering frequency and primary visual-dorsal default network coupling in the Parkinson’s disease hallucinators, relative to the non-hallucinating patients. This result confirmed that an increased propensity to mind-wander in the hallucinating patients was associated with greater connectivity between the default network and early visual regions. Controls showed a similar positive relationship between mind-wandering frequency and primary visual-dorsal default network coupling, however this association did not differ statistically from Parkinson’s disease non-hallucinators. Indeed, in healthy individuals, coupling between the default network and visual cortex has previously been associated with the degree of vivid detail experienced during mind-wandering episodes (Turnbull *et al.*, 2019). Considering the similar patterns between hallucinating patients and controls, this emphasises the overall interpretation of our results as one of a *preserved*, rather than *increased*, association between mind-wandering frequency and primary visual-default network in Parkinson’s disease hallucinations. However, unlike controls, this relative preservation is occurring in the context of impaired attention and primary visual dysfunction. When confronted with the resultant ambiguous sensory input, Parkinson’s disease hallucinators are therefore more likely to rely on top-down predictions to resolve a percept. This is consistent with the increased coupling between the default network and primary visual regions during recorded misperceptions in Parkinson’s disease hallucinators (Shine *et al.*, 2015b).

Our results show that, in the context of established Parkinson’s disease, patients with visual hallucinations exhibit greater mind-wandering frequency and a stronger association between mind-wandering frequency and primary visual-dorsal default network coupling, relative to non-hallucinators. However, when contrasted with healthy controls, Parkinson’s disease hallucinators were not statistically distinguishable on either of these variables. Non-hallucinators exhibited reduced mind-wandering compared to controls, accompanied by a negative association between mind-wandering frequency and primary visual-dorsal default network coupling. In light of the present evidence, we suggest two distinct trajectories in Parkinson’s disease: in Parkinson’s disease with hallucinations, relative to impaired sensory and attentional abilities, forms of internally generated mental imagery are relatively better preserved, which leads them to dominate perception – a process underpinned by increased primary visual-default network coupling; in Parkinson’s disease without hallucinations, such forms of cognition and the supporting neural circuity are compromised, consistent with a more generic course seen in ageing (Maillet and Schacter, 2016) and exacerbated in several neurodegenerative diseases (Geffen *et al.*, 2017; Gyurkovics *et al.*, 2018; O’Callaghan *et al.*, 2019).

Results from the post-hoc seed-to-voxel analysis confirmed a significant difference between the patient groups in the association between mind-wandering frequency and connectivity between the primary visual network and more restricted areas of the dorsal default network (posterior cingulate cortex, medial prefrontal cortex, right temporo-parietal junction and left inferior parietal lobule). This analysis enabled a more specific localisation of regions that might mediate the relationship between mind-wandering and primary visual network connectivity. Our findings implicated specific regions within the default network, including the medial prefrontal cortex, previously identified as a source of top-down influence over visual perception (Bar *et al.*, 2006; Summerfield *et al.*, 2006; Kveraga *et al.*, 2007). Previous work has also shown that stronger resting state coupling between the posterior cingulate / retrosplenial cortex and primary visual cortex correlates with the ability to actively imagine the future, which presumably relies upon scene construction processes (i.e., mentally envisaging a scene or event) (Villena-Gonzalez *et al.*, 2018). We also identified regions outside the default network, including the fusiform gyrus/inferior temporal gyrus, a higher-level region in the ventral visual processing stream; the inferior frontal gyrus, which has been identified (via effective connectivity) as a source of directed top-down influence over V1 during both mental imagery and perception (Dijkstra *et al.*, 2017); and the orbitofrontal cortex, a source of top-down predictions that refine object recognition during early visual processing (Bar, 2003; Bar *et al.*, 2006; Chaumon *et al.*, 2014). In the context of existing literature, the neural correlates identified in our analyses overlap with those that have been shown, in healthy people, to be important for coordinating internally generated predictions that influence both mental imagery and early visual perception.

Based on patterns of functional connectivity across the entire cerebral cortex, the default network is located on the opposite end of a principal connectivity gradient from brain regions supporting primary perceptual processing (e.g., the primary visual network) (Margulies *et al.*, 2016; Huntenburg *et al.*, 2018; Murphy *et al.*, 2018). Our results suggest that, in Parkinson’s disease with hallucinations, individual differences in mind-wandering frequency are associated with increasing loss of this inherent functional separation between the default network and primary visual areas. This is consistent with a previous task-based study, which found increased visual network coupling to the default network during misperceptions in Parkinson’s disease with hallucinations (Shine *et al.*, 2015b). A combination of increased internally generated thought and imagery, and increased influence from the default network over early visual regions, may therefore manifest in a neurocognitive endophenotype that is prone to hallucinate. Intriguingly, increased psychopathological features in the general population have been associated with elevated visual network-default network coupling (Elliott *et al.*, 2018), suggesting that the loss of functional segregation between these networks may play a role across neuropsychiatric disorders.

Given that a cardinal feature of mind wandering is its relative stimulus-independence (Mason *et al.*, 2007; Christoff *et al.*, 2016; Seli *et al.*, 2018), from a predictive processing framework this cognitive phenomenon would depend upon the activation (and possibly the exploration / finessing) of expectations or predictions (i.e., priors). Although the functional importance of mind-wandering remains unclear, it is possible that predictive model optimisation via the pruning of priors (i.e., reduction of model complexity or “overfitting”) during mind-wandering is analogous to predictive processing accounts for the functional importance of dreaming (Hobson and Friston, 2012), but occurs during the waking state with the associated benefits for adaptive fitness.

Previous work has shown that Parkinson’s disease patients who experience visual hallucinations are unable to contextually modulate sensory evidence accumulation (O’Callaghan *et al.*, 2017a). As the association between mind-wandering frequency and primary visual-dorsal default network coupling differentiated Parkinson’s disease patient groups with and without visual hallucinations, this pattern of network coupling may predispose the Parkinsonian brain to hallucinate by allowing relatively strong priors to dominate imprecise sensory evidence during the act of visual perception. Thus, while control subjects do not have significantly different mind-wandering frequencies or associations between mind-wandering frequency and primary visual-dorsal default network coupling compared with Parkinson’s disease hallucinators, they may not be prone to visual hallucinations due to intact mechanisms for the attribution of context-dependent precision estimation of sensory evidence. In keeping with this, previous work has suggested that a breakdown in the co-ordination of networks that support attention and saliency (including the dorsal and ventral attention networks) may be a crucial factor contributing to the inability to resolve imprecise sensory input in Parkinson’s disease hallucinations (Shine *et al.*, 2015a; 2015b). Our results highlight the importance of future studies that will directly examine the interplay between relatively preserved priors and imprecise sensory evidence in Parkinson’s disease hallucinations. One possible line of investigation is to use established tasks for conditioned hallucinations, which have been effective at identifying both relatively strong priors and neural regions supporting these priors in populations with auditory hallucinations (Powers *et al.*, 2016; 2017; Corlett *et al.*, 2018).

In summary, our study has identified mind-wandering frequency as a potential behavioural correlate of the pathophysiological default network, which has previously been implicated in the pathogenesis of visual hallucinations in Parkinson’s disease. To our knowledge, these findings provide the first evidence of a relationship between mind-wandering and visual hallucinations in a clinical population. Higher levels of mind-wandering indicate an increased propensity for detailed mental imagery, formation of contextual associations and spontaneous thought – all top-down processes that could furnish the content of visual hallucinations. Our results suggest a putative neural substrate to support excessive influence from these top-down processes over perception, by way of increased coupling between the default network and early visual regions. Elevated mind-wandering in association with loss of segregation between primary sensory and transmodal neural networks, therefore offers a specific neurocognitive marker for the top-down influences that may contribute to hallucinations across disorders.

## Acknowledgments

We thank Kelly Diederen for providing valuable comments on the manuscript. AJM is supported by an Australian Postgraduate Award through the University of Sydney. JMH is supported by a Western Sydney University Postgraduate Award. JAH is supported by a National Institutes of Aging Arizona Alzheimer’s Disease Core Center grant (P30 AG019610). MI is supported by an Australian Research Council Future Fellowship (FT160100096) and an Australian Research Council Discovery Project (DP180101548). SJGL is supported by an NHMRC-ARC Dementia Fellowship (#1110414). JMS is supported by a National Health and Medical Research Council CJ Martin Fellowship (1072403). CO is supported by a National Health and Medical Research Council Neil Hamilton Fairley Fellowship (1091310) and by the Wellcome Trust (200181/Z/15/Z). The study was supported by a Seed Grant from Parkinson’s NSW and funding to Forefront, a collaborative research group dedicated to the study of non-Alzheimer disease degenerative dementias, from the National Health and Medical Research Council of Australia program grant (#1037746 and #1095127).

## References

Adams RA, Stephan KE, Brown HR, Frith CD, Friston KJ. The computational anatomy of psychosis. Front. Psychiatry 2013; 4: 47.

Aminoff EM, Kveraga K, Bar M. The role of the parahippocampal cortex in cognition. Trends in Cognitive Sciences 2013; 17: 379–390.

Andrews-Hanna JR, Reidler JS, Sepulcre J, Poulin R, Buckner RL. Functional-anatomic fractionation of the brain’s default network. Neuron 2010; 65: 550–562.

Baldassano C, Chen J, Zadbood A, Pillow JW, Hasson U, Norman KA. Discovering Event Structure in Continuous Narrative Perception and Memory. Neuron 2017; 95: 709–721.e5.

Bar M, Kassam KS, Ghuman AS, Boshyan J, Schmid AM, Schmidt AM, et al. Top-down facilitation of visual recognition. Proceedings of the National Academy of Sciences 2006; 103: 449–454.

Bar M. A cortical mechanism for triggering top-down facilitation in visual object recognition. J Cogn Neurosci 2003; 15: 600–609.

Bates D, Mächler M, Bolker B, Walker S. Fitting Linear Mixed-Effects Models using lme4. 2014

Beck AT, Steer RA, Brown GK. Manual for the beck depression inventory-II. San Antonio, TX: Psychological Corporation 1996; 1: 82.

Bell AH, Summerfield C, Morin EL, Malecek NJ, Ungerleider LG. Encoding of Stimulus Probability in Macaque Inferior Temporal Cortex. Curr. Biol. 2016; 26: 2280–2290.

Buckner RL, Andrews-Hanna JR, Schacter DL. The brain’s default network: anatomy, function, and relevance to disease. Ann. N. Y. Acad. Sci. 2008; 1124: 1–38.

Cardinal RN, Aitken MRF. ANOVA for the Behavioral Sciences Researcher. 1st ed. Psychology Press; 2013.

Chaumon M, Kveraga K, Barrett LF, Bar M. Visual predictions in the orbitofrontal cortex rely on associative content. Cereb. Cortex 2014; 24: 2899–2907.

Christoff K, Irving ZC, Fox KCR, Spreng RN, Andrews-Hanna JR. Mind-wandering as spontaneous thought: a dynamic framework. Nat. Rev. Neurosci. 2016; 17: 718–731.

Collerton D, Perry E, McKeith I. Why people see things that are not there: A novel Perception and Attention Deficit model for recurrent complex visual hallucinations. Behav Brain Sci 2005; 28: 737–757.

Collerton D, Taylor J-P, Tsuda I, Fujii H, Shigetoshi N, Aihara K, et al. How Can We See Things That Are Not There? Current insights into complex visual hallucinations. Journal of Consciousness Studies 2016: 195–227.

Corlett PR, Horga G, Fletcher PC, Ben Alderson-Day, Schmack K, Powers AR III. Hallucinations and Strong Priors. Trends in Cognitive Sciences 2018: 1–14.

Dalrymple-Alford JC, MacAskill MR, Nakas CT, Livingston L, Graham C, Crucian GP, et al. The MoCA Well-suited screen for cognitive impairment in Parkinson disease. American Academy of Neurology 2010; 75: 1717–1725.

de Lange FP, Heilbron M, Kok P. How Do Expectations Shape Perception? Trends in Cognitive Sciences 2018; 22: 764–779.

Diederich NJ, Goetz CG, Stebbins GT. Repeated visual hallucinations in Parkinson’s disease as disturbed external/internal perceptions: focused review and a new integrative model. Mov Disord. 2005; 20: 130–140.

Dijkstra N, Zeidman P, Ondobaka S, van Gerven MAJ, Friston K. Distinct Top-down and Bottom-up Brain Connectivity During Visual Perception and Imagery. Scientific Reports 2017; 7: 5677.

Elliott ML, Romer A, Knodt AR, Hariri AR. A Connectome-wide Functional Signature of Transdiagnostic Risk for Mental Illness. Biol. Psychiatry 2018; 84: 452–459.

Ffytche DH, Creese B, Politis M, Chaudhuri KR, Weintraub D, Ballard C, et al. The psychosis spectrum in Parkinson disease. Nature Publishing Group 2017; 13: 81–95.

Fletcher PC, Frith CD. Perceiving is believing: a Bayesian approach to explaining the positive symptoms of schizophrenia. Nat. Rev. Neurosci. 2009; 10: 48–58.

Franciotti R, Delli Pizzi S, Perfetti B, Tartaro A, Bonanni L, Thomas A, et al. Default mode network links to visual hallucinations: A comparison between Parkinson’s disease and multiple system atrophy. Mov Disord. 2015; 30: 1237–1247.

Friston KJ. Hallucinations and perceptual inference. Behav Brain Sci 2005; 28: 764–766.

Geffen T, Thaler A, Gilam G, Simon EB, Sarid N, Gurevich T, et al. Reduced mind wandering in patients with Parkinson’s disease. Parkinsonism and Related Disorders 2017; 44: 38–43.

Gilbert CD, Li W. Top-down influences on visual processing. Nat. Rev. Neurosci. 2013; 14: 350–363.

Goetz CG, Tilley BC, Shaftman SR, Stebbins GT, Fahn S, Martínez-Martín P, et al. Movement Disorder Society-sponsored revision of the Unified Parkinson’s Disease Rating Scale (MDS-UPDRS): Scale presentation and clinimetric testing results. Mov Disord. 2008; 23: 2129–2170.

Gyurkovics M, Balota DA, Jackson JD. Mind-wandering in healthy aging and early stage Alzheimer’s disease. Neuropsychology 2018; 32: 89–101.

Hall JM, O’Callaghan C, Muller AJ, Ehgoetz Martens KA, Phillips JR, Moustafa AA, et al. Changes in structural network topology correlate with severity of hallucinatory behavior in Parkinson’s disease. Network Neuroscience 2019; 3: 521–538.

Hindy NC, Ng FY, Turk-Browne NB. Linking pattern completion in the hippocampus to predictive coding in visual cortex. Nature Neuroscience 2016; 19: 665–667.

Hobson JA, Friston KJ. Waking and dreaming consciousness: neurobiological and functional considerations. Progress in Neurobiology 2012; 98: 82–98.

Huntenburg JM, Bazin P-L, Margulies DS. Large-Scale Gradients in Human Cortical Organization. Trends in Cognitive Sciences 2018; 22: 21–31.

Irving ZC. Mind-wandering is unguided attention: accounting for the ‘purposeful’ wanderer. Philosophical Studies 2016; 173: 547–571.

Kok P. Perceptual Inference: A Matter of Predictions and Errors. Curr. Biol. 2016; 26: R809–11.

Kucyi A. Just a thought: How mind-wandering is represented in dynamic brain connectivity. NeuroImage 2017: 1–10.

Kveraga K, Boshyan J, Bar M. Magnocellular projections as the trigger of top-down facilitation in recognition. J. Neurosci. 2007; 27: 13232–13240.

Kveraga K, Ghuman AS, Kassam KS, Aminoff EA, Hämäläinen MS, Chaumon M, et al. Early onset of neural synchronization in the contextual associations network. Proc. Natl. Acad. Sci. U.S.A. 2011; 108: 3389–3394.

Maillet D, Schacter DL. From mind wandering to involuntary retrieval_ Age-related differences in spontaneous cognitive processes. Neuropsychologia 2016; 80: 142–156.

Margulies DS, Ghosh SS, Goulas A, Falkiewicz M, Huntenburg JM, Langs G, et al. Situating the default-mode network along a principal gradient of macroscale cortical organization. Proc. Natl. Acad. Sci. U.S.A. 2016; 113: 12574–12579.

Mason MF, Norton MI, Van Horn JD, Wegner DM, Grafton ST, Macrae CN. Wandering minds: the default network and stimulus-independent thought. Science 2007; 315: 393–395.

Murphy C, Jefferies E, Rueschemeyer S-A, Sormaz M, Wang H-T, Margulies DS, et al. Distant from input: Evidence of regions within the default mode network supporting perceptually-decoupled and conceptually-guided cognition. NeuroImage 2018; 171: 393–401.

O’Callaghan C, Hall JM, Tomassini A, Muller AJ, Walpola IC, Moustafa AA, et al. Visual Hallucinations Are Characterized by Impaired Sensory Evidence Accumulation: Insights From Hierarchical Drift Diffusion Modeling in Parkinson’s Disease. Biological Psychiatry: Cognitive Neuroscience and Neuroimaging 2017a; 2: 680–688.

O’Callaghan C, Kveraga K, Shine JM, Adams RB Jr, Bar M. Predictions penetrate perception: Converging insights from brain, behaviour and disorder. Consciousness and Cognition 2017b; 47: 63–74.

O’Callaghan C, Shine JM, Hodges JR, Andrews-Hanna JR, Irish M. Hippocampal atrophy and intrinsic brain network dysfunction relate to alterations in mind wandering in neurodegeneration. Proc. Natl. Acad. Sci. U.S.A. 2019; 116: 3316–3321.

O’Callaghan C, Shine JM, Lewis SJG, Andrews-Hanna JR, Irish M. Shaped by our thoughts--a new task to assess spontaneous cognition and its associated neural correlates in the default network. Brain and Cognition 2015; 93: 1–10.

Power JD, Barnes KA, Snyder AZ, Schlaggar BL, Petersen SE. Spurious but systematic correlations in functional connectivity MRI networks arise from subject motion. NeuroImage 2012; 59: 2142–2154.

Powers AR, Kelley M, Corlett PR. Hallucinations as top-down effects on perception. Biological Psychiatry: Cognitive Neuroscience and Neuroimaging 2016; 1: 393–400.

Powers AR, Mathys C, Corlett PR. Pavlovian conditioning-induced hallucinations result from overweighting of perceptual priors. Science 2017; 357: 596–600.

Schacter DL, Addis DR, Hassabis D, Martin VC, Spreng RN, Szpunar KK. The Future of Memory: Remembering, Imagining, and the Brain. Neuron 2012; 76: 677–694.

Seli P, Kane MJ, Smallwood J, Schacter DL, Maillet D, Schooler JW, et al. Mind-Wandering as a Natural Kind: A Family-Resemblances View. Trends in Cognitive Sciences 2018; 22: 479–490.

Shin D-J, Lee TY, Jung WH, Kim SN, Jang JH, Kwon JS. Away from home: the brain of the wandering mind as a model for schizophrenia. Schizophrenia Research 2015; 165: 83–89.

Shine JM, Keogh R, O’Callaghan C, Muller AJ, Lewis SJG, Pearson J. Imagine that: elevated sensory strength of mental imagery in individuals with Parkinson’s disease and visual hallucinations. Proceedings of the Royal Society B: Biological Sciences 2015a; 282: 20142047–20142047.

Shine JM, Muller AJ, O’Callaghan C, Hornberger M, Halliday GM, Lewis SJG. Abnormal connectivity between the default mode and the visual system underlies the manifestation of visual hallucinations in Parkinson’s disease: a task-based fMRI study. npj Parkinsons Disease 2015b; 1: 15003.

Shine JM, O’Callaghan C, Halliday GM, Lewis SJG. Tricks of the mind: Visual hallucinations as disorders of attention. Progress in Neurobiology 2014: 1–8.

Shirer WR, Ryali S, Rykhlevskaia E, Menon V, Greicius MD. Decoding subject-driven cognitive states with whole-brain connectivity patterns. Cereb. Cortex 2012; 22: 158–165.

Smallwood J, Schooler JW. The science of mind wandering: empirically navigating the stream of consciousness. Annu Rev Psychol 2015; 66: 487–518.

Spreng RN, Mar RA, Kim ASN. The Common Neural Basis of Autobiographical Memory, Prospection, Navigation, Theory of Mind, and the Default Mode: A Quantitative Meta-analysis. J Cogn Neurosci 2009; 21: 489–510.

Summerfield C, Egner T, Greene M, Koechlin E, Mangels J, Hirsch J. Predictive codes for forthcoming perception in the frontal cortex. Science 2006; 314: 1311–1314.

Turnbull A, Wang H-T, Schooler JW, Jefferies E, Margulies DS, Smallwood J. The ebb and flow of attention: Between-subject variation in intrinsic connectivity and cognition associated with the dynamics of ongoing experience. NeuroImage 2019; 185: 286–299.

Villena-Gonzalez M, Wang H-T, Sormaz M, Mollo G, Margulies DS, Jefferies EA, et al. Individual variation in the propensity for prospective thought is associated with functional integration between visual and retrosplenial cortex. CORTEX 2018; 99: 224–234.

Weil RS, Schrag AE, Warren JD, Crutch SJ, Lees AJ, Morris HR. Visual dysfunction in Parkinson’s disease. Brain 2016; 139: 2827–2843.

Weilnhammer VA, Stuke H, Sterzer P, Schmack K. The neural correlates of hierarchical predictions for perceptual decisions. Journal of Neuroscience 2018; 38: 5008–5021.

Whitfield-Gabrieli S, Nieto-Castañón A. Conn: a functional connectivity toolbox for correlated and anticorrelated brain networks. Brain Connectivity 2012; 2: 125–141.

Yao N, Shek-kwan Chang R, Cheung C, Pang S, Lau KK, Suckling J, et al. The default mode network is disrupted in parkinson’s disease with visual hallucinations. Hum. Brain Mapp. 2014; 35: 5658–5666.

Zabelina DL, Andrews-Hanna JR. Dynamic network interactions supporting internally-oriented cognition. Current Opinion in Neurobiology 2016; 40: 86–93.

