## Supplementary Material for "Mind-wandering in Parkinson’s disease hallucinations reflects primary visual and default network coupling"

**1. Detail for the scoring continuum of the thought sampling task and representative examples**

**2. Group comparison results for demographics, clinical characteristics and background neuropsychology measures**

**3. Effect of trial duration on scores achieved on the thought sampling task**
*(Figure S1)*

**4. Follow-up seed-to-voxel functional connectivity with a V1 seed in controls vs. Parkinson's disease non-hallucinators**
*(Figure S2)*

**1. Detail for the scoring continuum of the thought sampling task and representative examples**

Adapted from O’Callaghan et al. (O'Callaghan *et al.*, 2015). Reported thoughts were classified as follows:

**Level 1 = Stimulus-bound/Nothing**

Thoughts that are completely stimulus-bound, i.e., merely labelling the shape presented on the screen; and/or failure to generate any specific thoughts during stimulus presentation. Indicating no specific thoughts were generated or the individual’s thoughts were entirely stimulus-bound during the trial.

For example: “It was just a blue square, nothing else”; “Nothing”, “my mind was blank”

**Level 2 = Stimulus-/task-/environmental-dependent responses.**

Thoughts that are tied to stimulus attributes, task parameters, or environmental factors. These include, (i) Stimulus attributes: shape, colour; (ii) Task parameters: trial duration, purpose of task; (iii) Environment: external or sensory distractions

For example: (i) “That’s a nice round shape”; “The yellow is nice and bright”; “Yellow is my favourite colour”; (ii) “That one took a long time”; “I was wondering what the task was about”; “what is the point of this”; (iii) “It’s cold in this room”; “I could hear the birds outside”.

**Level 3 = Shape-related extrapolation**

Responses signifying an intermediate zone between stimulus-/task-dependent thinking and instances of “pure” mind wandering. Level 3 responses do not satisfy criteria for mind wandering as they still reflect a degree of reliance on the stimulus at hand. Thoughts have clearly shifted beyond the immediate perceptual features of the stimulus, yet the individual continues to rely upon perceptual features to generate some additional comparative mental imagery.

For example: “The yellow reminded me of the sun”; “The shape was like the one used by the Red Cross but it was the wrong colour”; “It reminds me of an opened umbrella”.

**Level 4 = Mind wandering**

Thoughts have moved beyond the immediate perceptual features of the shape/task and beyond comparisons that are tied to the stimulus. These responses are stimulus-/task-independent, and clearly demonstrate a move beyond stimulus attributes and comparisons tied to the stimulus, i.e., these thoughts do not directly relate to any properties of the stimulus, the task, or the environmental surroundings.

For example: "I thought about the people I saw today and how we chatted with them outside the unit"; "I thought of a sailing boat in the Greek Islands"; "I was wondering what time I'll be finished here and if I'll make it to the supermarket"; "I started thinking about what I will make for dinner tonight"; "I remembered being at school and trying to learn trigonometry".

### 2. Group comparison results for demographics, clinical characteristics and background neuropsychology measures

Parkinson's disease and control groups were matched for age and education [age: $F(2,75) = 1.79$, $p = 0.17$; education: $F(2,75) = 1.75$, $p = 0.18$]. PD+VH and PD-VH groups were matched on clinical variables, including global cognition [MoCA: $t(34.49) = -0.24$, $p = 0.81$], years of disease duration [$t(28.51) = -1.41$, $p = 0.17$], daily dopamine medication levels [DDE: $t(36) = -1.041$, $p = 0.30$], disease stage and motor severity [Hoehn and Yahr: $t(27.98) = -0.44$, $p = 0.66$; UPDRS III: $t(33.34) = -1.06$, $p = 0.30$], and depression levels [$t(26.81) = -1.28$, $p = 0.21$]. However, PD+VH performed worse than PD-VH on measures of attentional set-shifting and memory retention [TMT B-A: $t(21.75) = -2.87$, $p < 0.01$; Logical memory % retention: $t(27.67) = 2.395$, $p < 0.05$]. Working memory performance did not differ between the patient groups [Digit span backward: $t(32.44) = -0.73$, $p = 0.47$]. See Table 1 in main manuscript.

#### 3. Effect of trial duration on scores achieved on the thought sampling task

***All groups showed an increased tendency to mind wander on longer trials***

A prediction of the task is that performance will be modulated by the duration of trials, i.e., that higher scores would occur on longer trials. This prediction was confirmed in the current study. There was a main effect of trial duration [F(2,150) = 12.46, $p < 0.0001$]. Post hoc t-tests showed that, regardless of group, higher average scores per trial were obtained in the long trials compared to the short trials ($p < 0.05$). The main effect of group was significant [F(2,75) = 3.97, $p < 0.05$], with post-hoc tests confirming that PD-VH had lower average trial scores compared to PD+VH and controls ($p < .05$). The group x duration interaction was not significant [F(4,150) = 0.61, $p = 0.65$].

Below is a plot of the average score achieved (ranging from Levels 1- 4) for each of the trial durations: Short: ≤20 seconds, Medium: 30-60 seconds, Long: ≥90 seconds. All groups had higher average scores on the Medium and Long trials, compared to the Short trials.

**Figure S1**

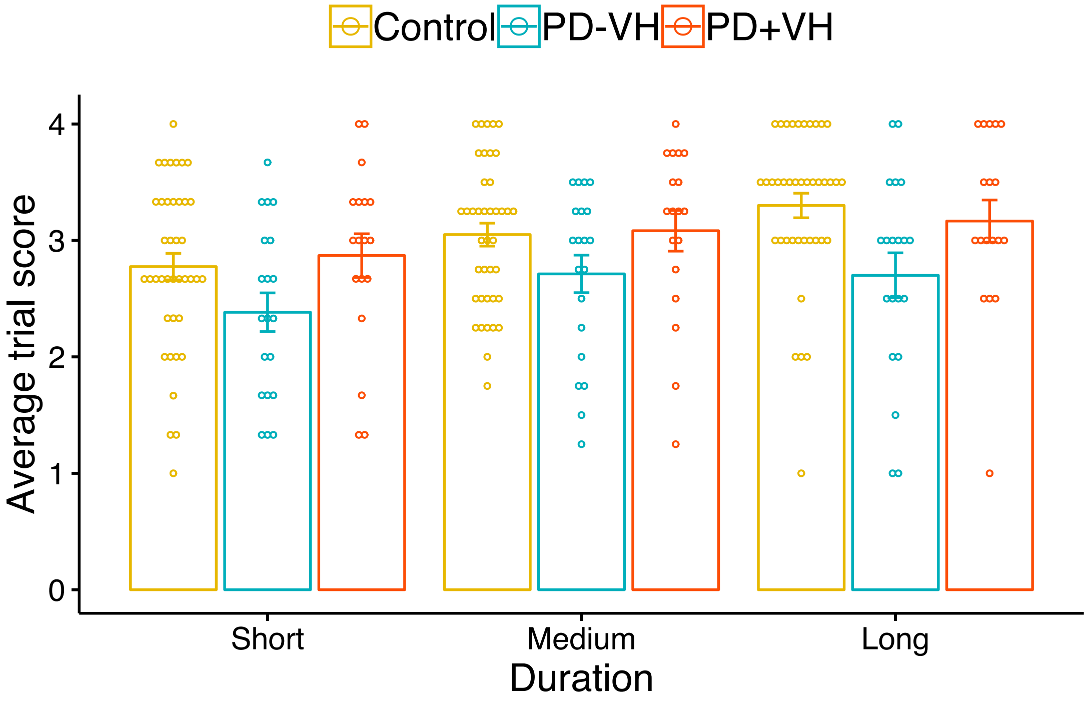

**4. Follow-up seed-to-voxel functional connectivity with a V1 seed in controls vs. PD-VH**

Seed-to-voxel connectivity between V1 seed and whole brain in PD+VH vs. PD-VH.

**Figure S2**

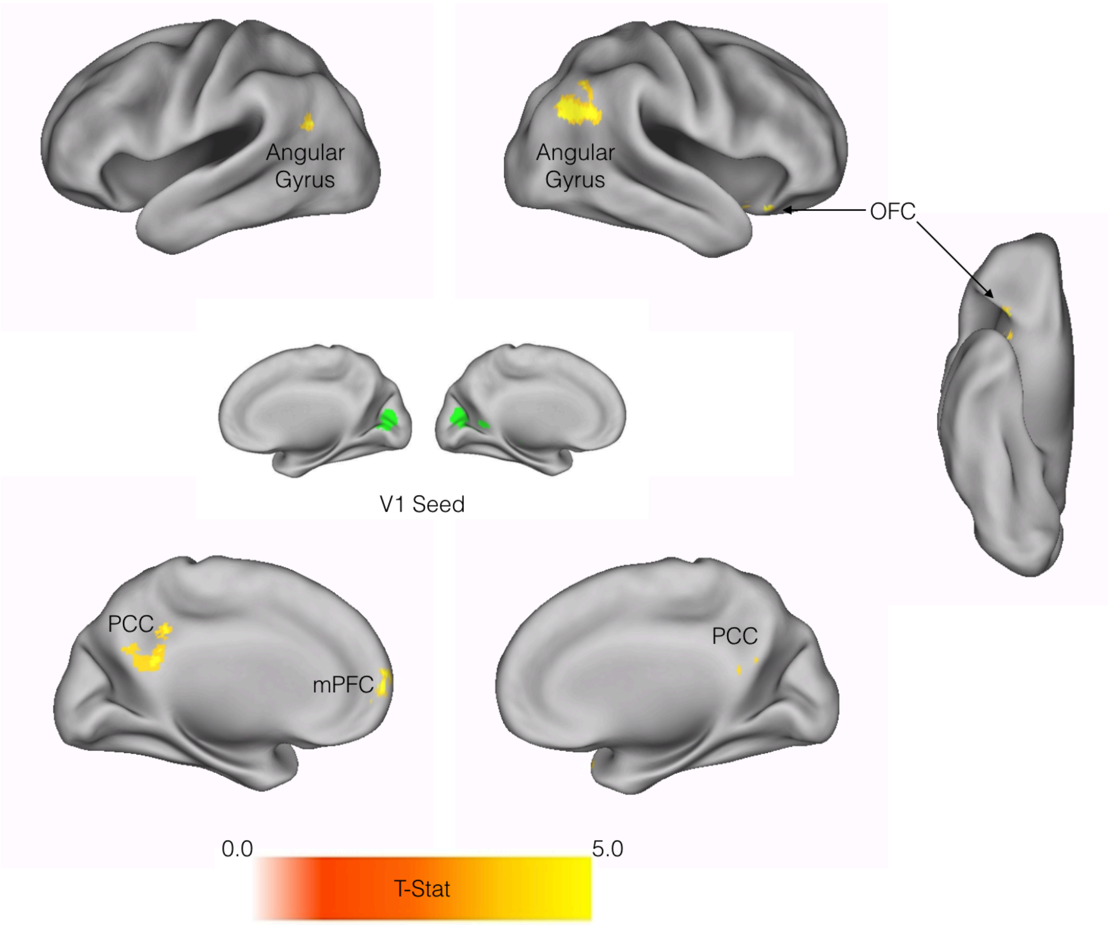


Table S1. Peak coordinates from seed-to-voxel analysis Controls vs PD-VH

| Brain Region | x | y | z | Voxels |
|---|---|---|---|---|
| R Oribitofrontal Cortex | 36 | 20 | -20 | 135 |
| L Medial Prefrontal Cortex | -10 | 60 | 00 | 118 |
| L Posterior Cingulate Cortex | 00 | -50 | 20 | 305 |
| R Angular Gyrus | 50 | -64 | 32 | 224 |
| L Angular Gyrus | -42 | 50 | 16 | 151 |

### References

O'Callaghan C, Shine JM, Lewis SJG, Andrews-Hanna JR, Irish M. Shaped by our thoughts – A new task to assess spontaneous cognition and its associated neural correlates in the default network. Brain and Cognition 2015; 93(0): 1-10.
